# DropletQC: improved identification of empty droplets and damaged cells in single-cell RNA-seq data

**DOI:** 10.1101/2021.08.02.454717

**Authors:** Walter Muskovic, Joseph E Powell

**Author notes:** Corresponding author; Joseph E Powell –. Walter Muskovic –.

## Abstract

Advances in droplet-based single cell RNA-sequencing (scRNA-seq) have dramatically increased throughput, allowing tens of thousands of cells to be routinely sequenced in a single experiment. In addition to cells, droplets capture cell-free “ambient” RNA predominately caused by lysis of cells during sample preparation. Samples with high ambient RNA concentration can create challenges in accurately distinguishing cell-containing droplets and droplets containing ambient RNA. Current methods to separate these groups often retain a significant number of droplets that do not contain cells – so called empty droplets. Additional to the challenge of identifying empty drops, there are currently no methods available to detect droplets containing damaged cells, which comprise of partially lysed cells – the original source of the ambient RNA. Here we describe *DropletQC*, a new method that is able to detect empty droplets, damaged, and intact cells, and accurately distinguish from one another. This approach is based on a novel quality control metric, the nuclear fraction, which quantifies for each droplet the fraction of RNA originating from unspliced, nuclear pre-mRNA. We demonstrate how *DropletQC* provides a powerful extension to existing computational methods for identifying empty droplets such as *EmptyDrops*. We have implemented *DropletQC* as an *R* package, which can be easily integrated into existing single cell analysis workflows.

## Main text

Droplet-based single cell RNA-sequencing (scRNA-seq) methods utilise microfluidics to encapsulate individual cells in nanolitre droplet emulsions, a technique that has dramatically increased throughput compared to plate-based protocols[1]. While encapsulating cells, droplets also capture cell-free ambient RNA, a complex mixture of transcripts released from damaged, stressed, and dying cells, often exacerbated during dissociation of solid tissues. This ambient RNA creates challenges for downstream analyses and the biological interpretation of results as most analysis methods are based on the assumption that a droplet contains RNA from a single cell. To combat this problem, several methods have been developed to estimate and remove its contribution to gene expression [2–4].

High levels of ambient RNA also create challenges in accurately identifying cell-containing droplets. This is a particular problem for data generated from solid tissues, where more fragile cells are more likely to become damaged during dissociation, and contribute to ambient RNA. We thus have three scenarios that need to be differentiated: empty droplets containing high concentrations of ambient RNA; droplets containing damaged cells; and droplets containing cells with limited ambient RNA. Using cut-offs based on the total number of RNA fragments assigned to each droplet, such as those originally proposed by Macosko *et al*. [5] and Zheng *et al*. [6], risks both including empty droplets and excluding small cells with below-average RNA content. The *EmptyDrops* method [7] addresses this issue through a more sophisticated approach, calculating the profile of the ambient RNA pool and testing each barcode for significant deviations from this profile. A favoured alternative to simple UMI cut-offs, this method has been integrated as the default cell-calling algorithm in the widely used *CellRanger* pipeline [6]. However, cell-free droplets with high ambient RNA concentration and damaged cells are still retained by this method.

Here, we present *DropletQC*, a new method that is able to simultaneously improve the detection of cell free droplets and droplets containing damaged cells. Taking advantage of the observation that unspliced and spliced mRNAs can be distinguished in common scRNA-seq protocols [8], we develop a novel metric: the nuclear fraction. The nuclear fraction quantifies, for each droplet, the proportion of RNA originating from unspliced pre-mRNA. Ambient RNA consists predominantly of mature cytoplasmic mRNA. This may arise as RNA is released from damaged cells in which the nuclear envelope remains intact, or capped and polyadenylated transcripts may be more stable in the extracellular environment (**Figure 1**). Regardless, droplets that contain only ambient RNA have a low nuclear fraction compared to droplets containing cells. In contrast, damaged cells due to the depletion of cytoplasmic RNA, will have a higher nuclear fraction compared to intact cells. By using the nuclear fraction score in combination with UMIs per droplet, we are able to accurately distinguish between empty droplets, damaged cells, and intact cells.

**Figure 1.**
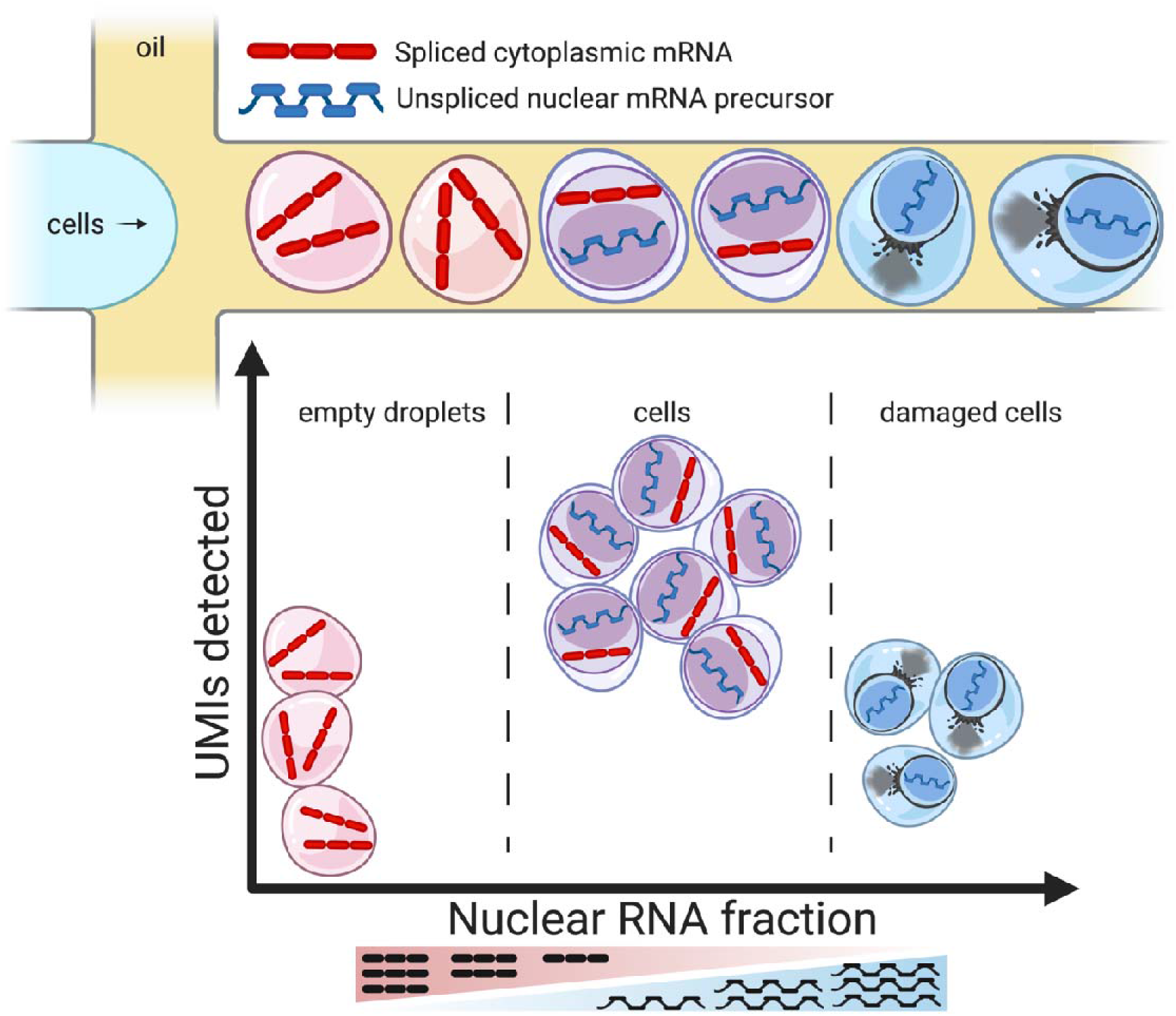
Illustration of how the nuclear fraction, in combination with the library size of each droplet, can be used to separate the populations of empty droplets, intact cells and damaged cells.

To assess the ability of *DropletQC* to identify both empty droplets and droplets containing damaged cells, we applied it to four independent scRNA-seq datasets; embryonic mouse brain, glioblastoma tumour, peripheral blood mononuclear cells (PBMCs), and Hodgkin’s lymphoma tumour. To determine whether *DropletQC* could identify empty droplets missed by current methods, all barcodes were first filtered using *EmptyDrops*, as implemented in *DropletUtils* [7]. *DropletQC* identified an additional 9.5% of mouse brain, 6.0% of Hodgkin’s lymphoma, 4.0% of glioblastoma and 0.4% of PBMCs as empty droplets (**Figure 2**, **Table S1**). Cells from dissociated tissue (**Figure 2a-c**) contained more empty droplets with high RNA content than PBMCs (**Figure 2d**), suggesting ambient RNA may be released from cells damaged during sample preparation.

**Figure 2.**
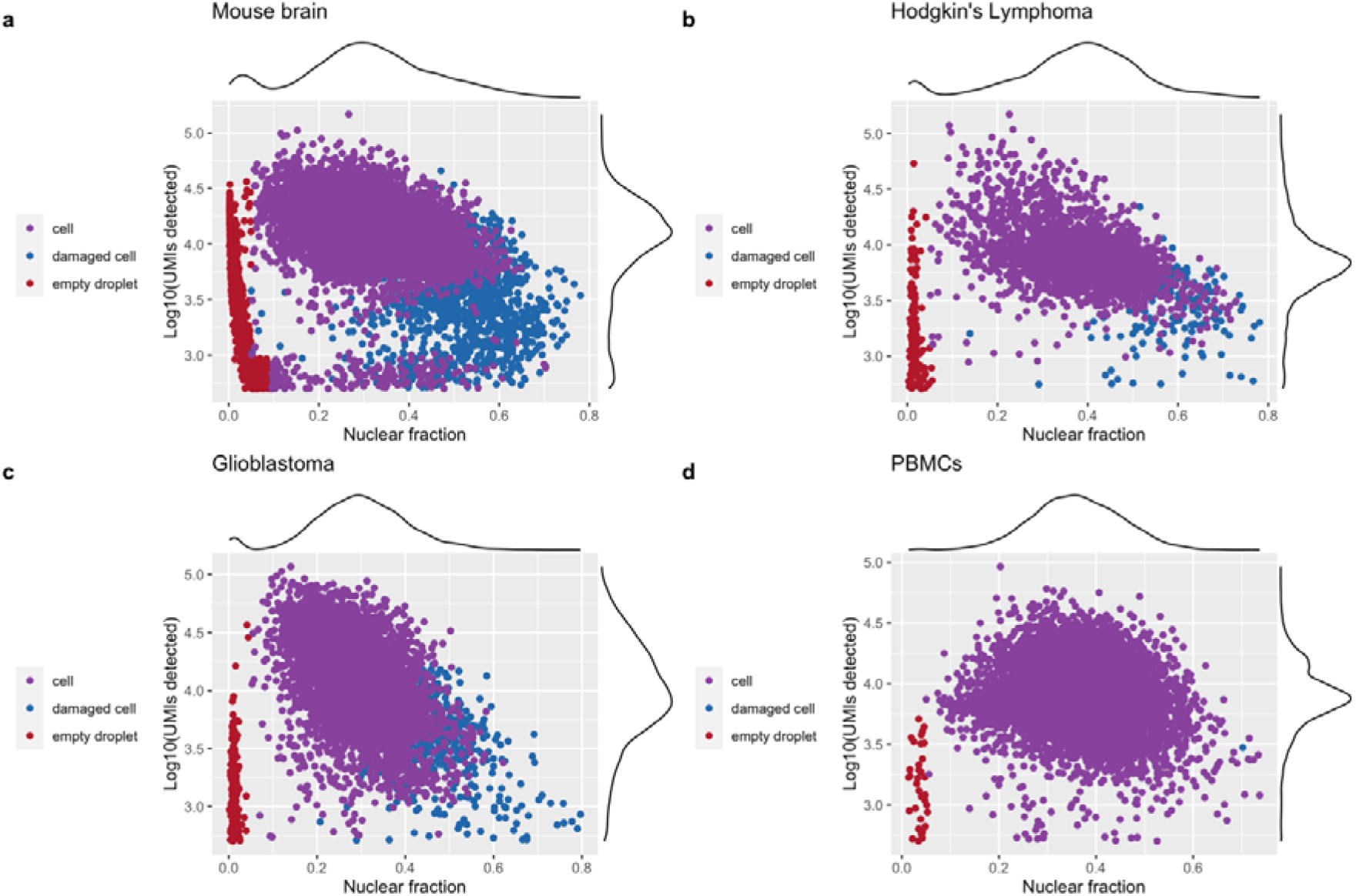
DropletQC identifies empty droplets and damaged cells in four heterogeneous scRNA-seq datasets. Total UMI counts (y-axis) and nuclear fraction scores (x-axis) are shown for each cell, with colours representing the status of each cell assigned by DropletQC. Empty droplets contain less RNA than cells and a higher fraction of cytoplasmic RNA (low nuclear fraction score). Damaged cells contain less RNA than intact cells and a higher proportion of unspliced RNA fragments (high nuclear fraction score).

Following identification of empty droplets, droplets containing damaged cells are identified using expectation maximisation and a Gaussian mixture model to separate them from droplets containing intact cells. As both the total UMI count and nuclear fraction scores display distinct distributions for different cell types (**Figure S1**), it is necessary to first group cells by type. Cells were annotated for each sample using a combination of gene markers and supervised classification with *scPred* [9]. Of the remaining cells, 14.0% of mouse brain, 5.2% of Hodgkin’s lymphoma, 9.8% of glioblastoma tumour cells and one PBMC cell were identified as damaged cells (**Table S1**).

As an additional test of the ability of *DropletQC* to identify damaged cells, we applied the method to data from a recent investigation on the effects of cryopreservation on the transcriptomes of macaque microglia [10]. *DropletQC* revealed an increase in the proportion of damaged cells following cryopreservation from 4.1% to 13.8% (**Figure S2, Table S2**). These findings have implications for the suitability of prospectively archiving samples for scRNA-seq studies through cryopreservation and demonstrates the utility of *DropletQC* for similar studies.

By default, the *DropletQC* software provides a flag for empty droplets and damaged cells, but does not automatically remove them from the dataset. Depending on the biological analyses, damaged cells may retain useful information, and as such it may be desirable to retain this metadata throughout downstream analyses. Similarly, cells such as erythrocytes, which contain small amounts of mature mRNA, may be misidentified as empty droplets and can be rescued downstream if desired.

For samples with large percentages of ambient RNA, some damaged cells and empty droplets may be missed by *DropletQC*. However, these can be identified by their low RNA content (**Figure 2a**) and may be easily flagged using a minimum UMI threshold. Calculation of the nuclear fraction, identification of empty droplets and damaged cells are implemented as separate functions within the *DropletQC* package. In summary, we have shown that *DropletQC* is able to successfully identify both empty droplets and damaged cells in data from a range of tissue types.

## Methods

### Nuclear fraction calculation

The *DropletQC* method first calculates the nuclear fraction for each droplet, which is the proportion of RNA fragments that originate from intronic regions. It is calculated as:

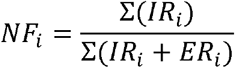

Where *NF_i_* is the nuclear fraction for droplet *i*, *IR_i_*, are the reads that map to intronic regions for droplet *i*, and *ER_i_* are those that map to exonic regions. We have implemented two methods to map reads to either intronic or exonic regions. The first, takes advantage of region tags, such as those added by 10x Genomics’ Cell Ranger count analysis pipeline that identify the region type of each genome-aligned RNA fragment; exonic, intronic or intergenic. These are efficiently counted using the nuclear_fraction_tags function to calculate a nuclear fraction score for each provided cell barcode. Alternatively, if region tags are missing, our second method assesses aligned reads for overlap with intronic regions using the nuclear_fraction_annotation function in combination with a user-provided gene annotation file. To speed up processing of indexed, coordinate-sorted alignment files, reads are split across a user-specified number of genomic regions to allow parallel computation. The four samples presented in the manuscript were processed with 8 CPUs and 16Gb of RAM with an average processing time of 106 seconds per 100 million reads using the nuclear_fraction_tags function and 132 seconds per 100 million reads using the nuclear_fraction_annotation function.

### Identifying empty droplets and damaged cells

Empty droplets are classified as all barcodes that fall below a defined nuclear fraction threshold. To identify a suitable threshold, a kernel density estimate is calculated using the nuclear fraction scores. The first derivative of the estimate is then calculated to identify the local minimum immediately following the first peak, corresponding to the population of empty droplets. If the automatically selected cut-off misidentifies the empty droplet population, two user-adjustable parameters are provided; a nuclear fraction threshold and a total UMI threshold, above which all barcodes are marked as cells.

To identify droplets containing damaged cells, barcodes are assessed separately for each cell type. It is assumed damaged cells have both a lower UMI count and higher nuclear fraction score than undamaged cells. We therefore use a two component (*k*) gaussian mixture model to classify droplets containing damaged cells:

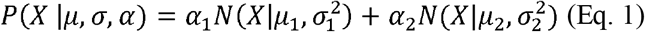

Where *X* is a dataset with *log*_10_(*UMI*) and estimated nuclear fractions for *1-n* droplets of a given cell type. *μ* and *σ^2^* are the mean and variance, and *α* represents the mixing weight of a given component. The initial model parameters are calculated as:

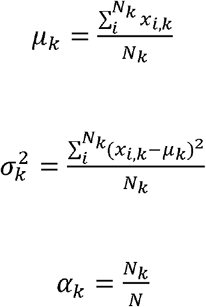

Where *N_k_* is the number of data points in the *k^th^* component. Following the initialisation, we estimate parameters using expectation maximisation by asking what is the posterior probability that a droplet (*x_i_*) belongs to component *k_j_*:

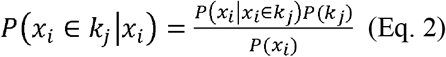

Where,

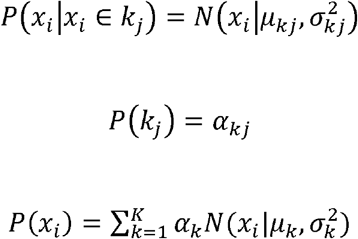

*N_k_* in the initial component parameters are replaced with the posterior probability and recalculated, with these steps repeated until convergence determined using the Bayesian information criterion. This model identifies the minimum separation required between the identified distributions for a population of droplets to be marked as damaged. We then label droplets as containing a damaged cell based on: a higher mean nuclear fraction and lower mean UMI than the cell population; a mean nuclear fraction greater than the cell population mean by a user-adjustable amount (default 0.15); a mean UMI count lower than the cell population (default 50%).

### Data

#### Cell filtering and annotation

Prior to calculating the nuclear fraction score, all cell barcodes were assessed for a significant deviation from the ambient RNA expression pattern using the *EmptyDrops* method implemented in the *DropletUtils* [7]. The lower bound on the total UMI count used to identify empty droplets was increased from 100 to 500 and all other parameters were left at their default values. Barcodes below a false discovery rate threshold of 1% were excluded. Remaining barcodes were additionally filtered for a maximum mitochondrial gene content of 15% to exclude low quality cells. Mouse brain and PBMC cell types were annotated by supervised classification with the *scPred* [9] using annotated PBMC [11], mouse brain [12] and developing mouse brain [13] reference datasets. The glioblastoma sample cell types were identified using cell-type specific gene markers for oligodendrocytes (*MAG, MOG, MBP*), microglia/macrophages (*C1QA, AIF1, LAPTM5*), T cells (*CD2, CD3D, CD3E*) and endothelial cells (*CD34, ESAM, APOLD1*) [14–17]. Hodgkin’s lymphoma cell types were classified using marker genes for B cells (*MS4A1*), macrophages (*CD68, IDO1*), plasmacytoid dendritic cells (*CLEC4C, NRP1*), erythrocytes (*HBB, HBA1, HBA2*), cytotoxic T cells (*GZMA, GZMK, IFNG*), regulatory T cells (*FOXP3, IL2RA, IKZF2*), T helper cells (*CXCL13, PDCD1, FABP5*), naïve T cells (*CCR7, IL7R, LEFT*), progenitor (*CD34*) and mast cells (*TPSAB1, TPSB2, KIT*) [18,19].

#### Availability of data and materials

The four single-cell gene expression datasets presented in the manuscript are made publicly available through the 10x Genomics website: https://support.10xgenomics.com/single-cell-gene-expression/datasets. The macaque microglia expression data is available from the NCBI GEO database, under accession GSE162663. All of the code used to produce the analyses and figures presented in the manuscript, along with links to individual datasets, are available through GitHub at https://github.com/powellgenomicslab/dropletQC_paper. *DropletQC* is available as an *R* package at https://github.com/powellgenomicslab/DropletQC.

## Supplementary Figures

**Supplementary Figure 1.**
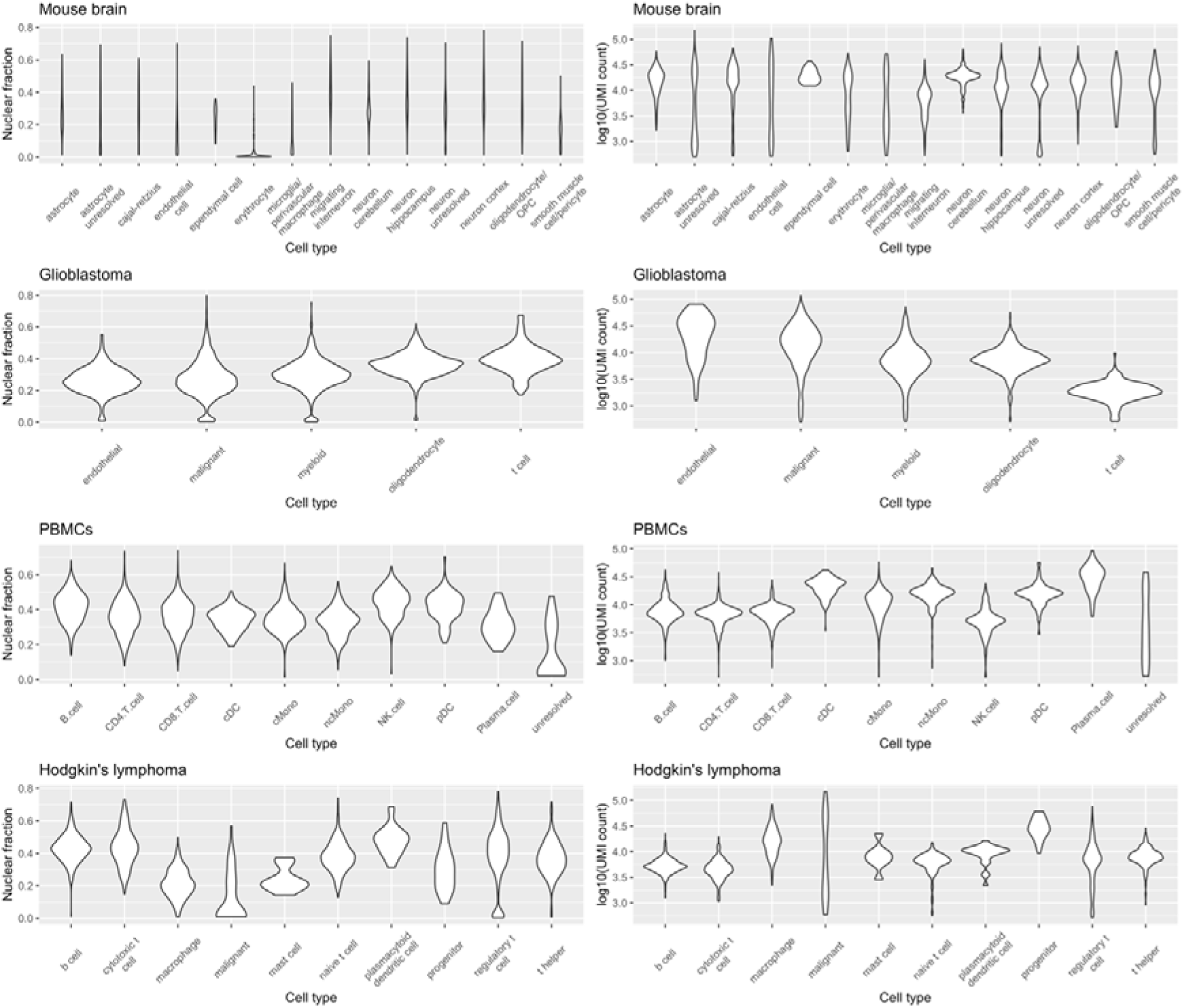
Different cell types have distinct distributions of nuclear fraction scores and UMI counts.

**Supplementary Figure 2.**
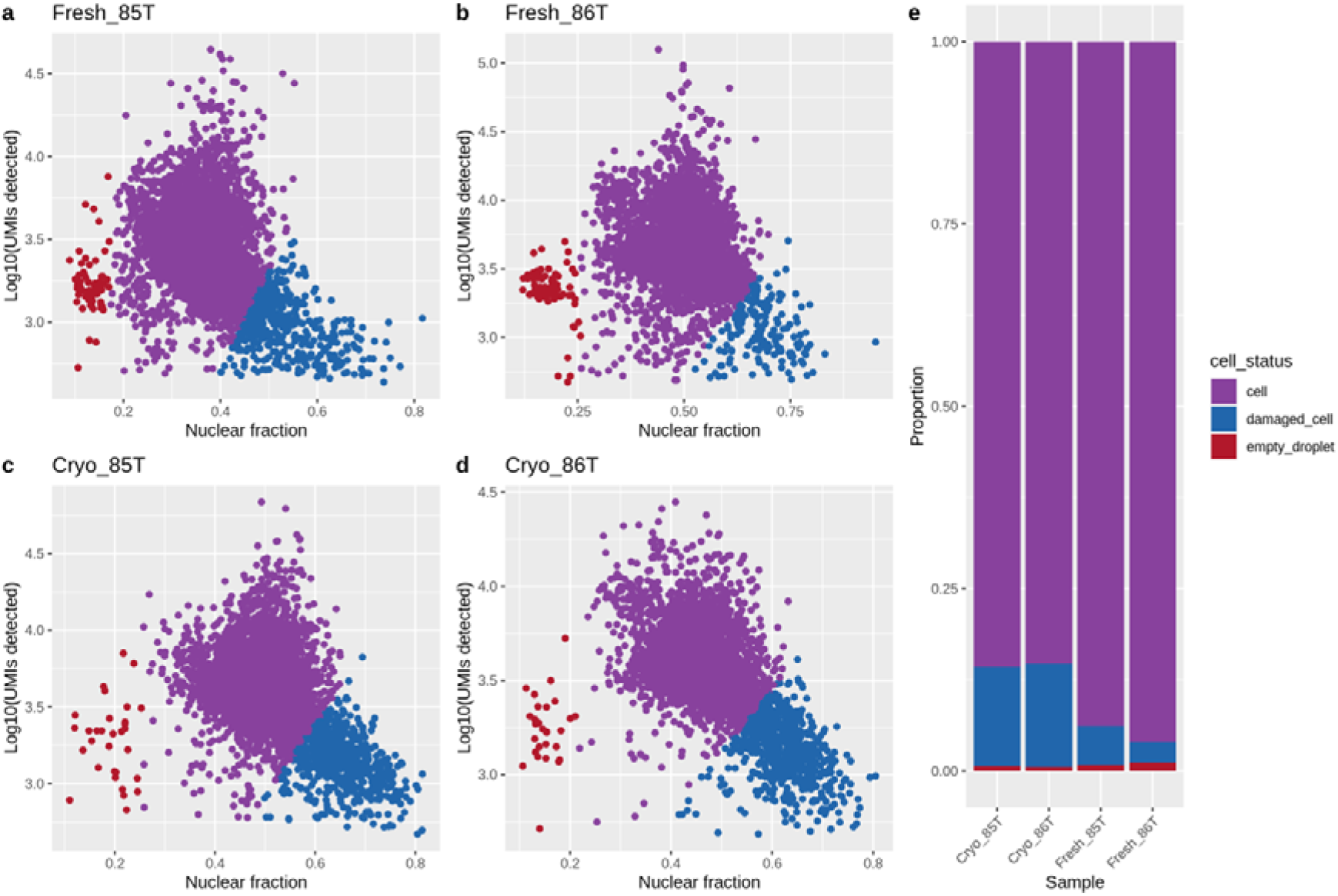
DropletQC identifies an increased proportion of damaged cells in cryopreserved microglia samples. (c-d) compared to fresh tissue samples (a-b). Total UMI counts (y-axis) and nuclear fraction scores (x-axis) are shown for each cell, with colours representing the status of each cell assigned by DropletQC. The stacked bar chart (e) illustrates the proportion of empty droplets and damaged cells for each sample.

## Supplementary Tables

**Supplementary Table 1.** Summary of the percentage of empty droplets and damaged cells identified in four heterogeneous scRNA-seq datasets after filtering with EmptyDrops and a maximum mitochondrial gene content of 15%.

| Sample | Empty droplets | Damaged cells |
| --- | --- | --- |
| Embryonic mouse brain | 9.53% | 14.00% |
| Glioblastoma tumour | 3.99% | 9.77% |
| Peripheral blood mononuclear cells | 0.37% | 0.01% |
| Hodgkin's lymphoma tumour | 6.04% | 5.16% |

**Supplementary Table 2.** Summary of the percentage of empty droplets and damaged cells identified in four macaque microglia scRNA-seq datasets.

| Sample | Empty droplets | Damaged cells |
| --- | --- | --- |
| Fresh_85T | 0.80% | 5.40% |
| Fresh_86T | 1.15% | 2.84% |
| Cryo_85T | 0.64% | 13.40% |
| Cryo_86T | 0.57% | 14.10% |

